# A vascular chip for disease-relevant flow shear stress topology

**DOI:** 10.64898/2026.07.07.736911

**Authors:** Kexin Li, Shiyi Yang, Ke Hu, Zhuqing Liang, Xing Zhang, Junjie Yang, Umberto Morbiducci, Valentina Mazzi, Diego Gallo, Li Wang, Min Wang, Xiaoning Sun, Zengsheng Chen, Anqiang Sun, Lingqian Chang, Yundai Chen, Yuehong Zheng, Xiao Liu

## Abstract

Vascular chips have advanced endothelial mechanobiology by enabling controlled responses to hemodynamic cues, yet disease-relevant wall shear stress (WSS) modeling remains limited. Simplified one-dimensional flow shear systems, designed mainly for physiological mechanobiology, miss the topological organization of pathological flow, whereas patient-specific vascular models capture complex hemodynamics but sacrifice generality and imaging compatibility. Here we develop a programmable vascular chip that converts disease-associated WSS topology into a physiologically parameterized experimental input. The device reconstructs a representative pathological shear-topology field on endothelial layer, supports stationary and physiologically paced oscillatory flow modes, and integrates matched unidirectional-shear references within the same chip. Using this system, we show that oscillatory WSS topology destabilizes endothelial monolayers, drives asymmetric collective emergent behaviors, impairs actin–nuclear mechanotransduction, accompanied by nuclear softening and enhanced perinuclear nanoparticle uptake. Integrated live-cell imaging, fluorescence analysis, Brillouin microscopy, and transport assays enable multimodal phenotyping across collective, subcellular mechanical and functional scales. By making disease-relevant WSS topology experimentally controllable, this vascular-chip framework bridges computational hemodynamics and experimental mechanomedicine, supporting standardized vascular disease modeling and functional screening.

## Introduction

Microphysiological systems and organ-on-chip technologies have become powerful platforms for modeling disease in controllable, human-relevant formats^1–5^. For vascular pathologies, the translational value of a vascular chip depends on whether it can reconstruct the hemodynamic microenvironment that shapes endothelial dysfunction, lesion formation and therapeutic response^6–10^. Endothelial cells lining the vessel wall are continuously exposed to wall shear stress (WSS), a mechanical cue that regulates endothelial homeostasis, inflammatory activation, barrier integrity, and molecular transport^11–13^. As vascular-chip design moves from reproducing physiological perfusion toward modeling disease-specific mechanics, the fidelity of WSS reconstruction has become a central benchmark of disease relevance.

Current approaches to vascular WSS modeling face a persistent trade-off between experimental generality and pathological fidelity. Straight-channel and parallel-plate systems precisely control shear magnitude and waveform and have provided the foundation for vascular mechanobiology^14,15^. More advanced vessel-on-chip designs can introduce curved channels, step flows, vascular architectures or local disturbed-flow regions, but they still largely represent WSS through scalar magnitude, temporal waveform or simplified flow direction^16–18^. Patient-specific three-dimensional vascular models offer substantial advantages for capturing anatomy-dependent hemodynamics, but face challenges such as difficulties to standardization, high-throughput screening, live imaging and quantitative phenotyping at cellular resolution^19^. Thus, existing platforms are either experimentally scalable but mechanically oversimplified, or anatomically realistic but difficult to deploy as generalizable vascular disease models.

The spatial topology of the WSS vector field is closely related to many vascular diseases and should be constructed in generalizable models as a general disease-relevant mechanical signature. In lesion-prone regions such as arterial bifurcations, stenoses and aneurysms, WSS is organized as a vector field with conserved topological features, including fixed points where local shear vectors vanish and neighboring vectors converge and diverge along defined directions^20,21^. WSS topological skeletons have been linked to impaired near-wall transport, atherogenic remodeling, plaque-prone regions and restenosis risk, with information that can be independent of conventional low-WSS metrics^22–25^.

The disease-relevant vascular chip becomes increasingly important amid the rise of mechanomedicine, an emerging paradigm that frames disease-associated mechanical signatures as measurable biomarkers, causal drivers and therapeutic targets, requiring platforms that can connect pathological mechanics to functional outcomes across scales^26–29^. For vascular disease modeling, this means that a chip should not only generate controlled flow, but also reproduce disease-relevant WSS topology and quantify its consequences for collective emergent behaviors^30,31^, actin-nuclear mechanotransduction and transport-related function^32–34^. Such transport-related phenotypes are particularly relevant to vascular-targeted nanotherapeutics, because endothelial barrier state, intracellular trafficking and molecular uptake directly influence therapeutic access at diseased vascular sites^12,35,36^.

Here we present a vascular chip platform that translates disease-associated WSS topological fields into physiologically parameterized experimental inputs, supporting both steady and cardiac-cycle oscillatory flow modes and incorporating matched unidirectional shear controls within a single device. Using this system, we show that oscillatory WSS topology regulates endothelial collective organization, actin–nuclear mechanotransduction and nanoparticle transport under matched shear conditions. By combining topological flow programming with multimodal phenotyping, this work provides a generalizable vascular-chip framework for disease-relevant modeling, mechanomedicine and functional screening.

## Results

### Vascular-chip design for disease-relevant flow shear stress topology

We developed a cross-junction vascular chip to engineer WSS topology as a controllable experimental variable. The design focuses on a saddle-type WSS pattern, a conserved feature of lesion-prone vascular geometries in which shear vectors vanish at a central fixed point, forming a characteristic convergent–divergent structure. (Fig. 1a). In active matter language, such saddle point topology is structurally analogous to a −1 topological defect, in the sense that both share the same integer index (winding/Poincaré index = −1)^24,37^. We therefore refer to the reconstructed shear field as −1 topological shear stress (−1 TSS) throughout this study. Rather than replicating the full three-dimensional anatomy of diseased vessels, the chip extracts this minimal, disease-defining topological feature and applies it as a defined mechanical input to endothelial monolayers.

**Fig. 1.**
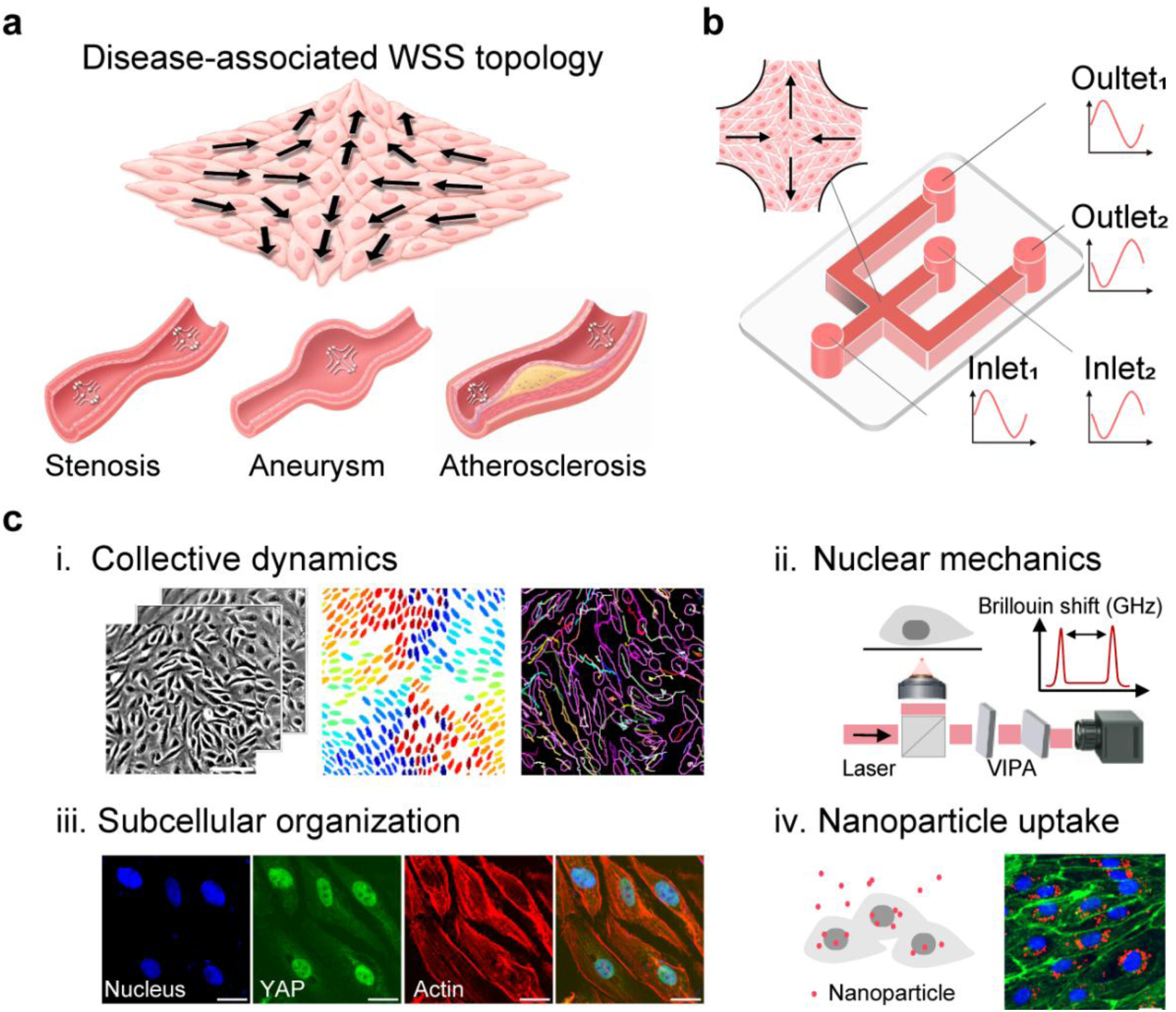
Vascular-chip design for disease-relevant flow shear stress topology. **a**, Disease-associated wall shear stress (WSS) topology represented by a −1 defect topological shear stress (TSS) field. The field exhibits orthogonal axes of vector convergence and divergence, illustrating a representative two-dimensional shear-topology pattern. The lower schematics show stenotic, aneurysmal and atherosclerotic vessels as examples of pathological vascular geometries associated with disturbed WSS patterns. **b**, Cross-junction vascular chip in which regulated inlet and outlet flows establish a -1 TSS field at the junction. **c**, Overview of chip-compatible analyses used to characterize endothelial responses across scales, including i, collective dynamics; ii, nuclear mechanics; iii, subcellular organization; and iv, nanoparticle uptake. Scale bars, 200 μm in c-i and 20 μm in c-iii and c-iv.

The device is built around a symmetric cross-junction geometry with four independently controlled flow channels (Fig. 1b). Under balanced steady perfusion fluid enters from one opposing channel pair and exits through the perpendicular pair, creating a saddle point at the junction center, where the local WSS vector approaches zero and the surrounding shear vectors converge along the inflow axis and diverge along the outflow axis. This mode generates stationary topological shear stress (STSS). By prescribing phase-shifted sinusoidal flow waveforms to opposing channels, the topological core can be driven into controlled periodic oscillation along the principal axis to generate oscillatory TSS (OTSS), while maintaining matched local shear levels. Straight-channel segments within the same device provide matched unidirectional shear stress (USS) references, enabling internally controlled comparisons between topological and non-topological shear environments while minimizing cross-device variability.

This programmable chip provides a unified platform for multiscale analysis (Fig. 1c). The same system enables live imaging of endothelial collective dynamics, label-free Brillouin assessment of nuclear mechanics, fluorescence-based characterization of cytoskeletal and nuclear organization, and nanoparticle uptake assays for transport function. By performing these measurements under a shared topological mechanical input, the platform links −1 TSS-induced responses from monolayer-scale organization to subcellular mechanotransduction and vascular-delivery-relevant transport behavior.

### Control and validation of topological flow shear stress in the vascular chip

To characterize the mechanical stimuli exerted by blood flow on endothelial layers, we simulated pulsatile flow in a patient-specific stenotic coronary artery. Analyzing the WSS vector field on the luminal surface, we identified saddle fixed points both upstream and downstream of the stenosis (Fig. 2a, Fig. S1a and Table S1), consistent with the preferential locations of atherosclerosis rupture and thrombus formation, respectively^6,22,23,38^. Time-resolved analysis further showed that the TSS pattern underwent lateral oscillatory displacement along the vessel wall during the cardiac cycle (Fig. S1b, c and Supplementary Video 1). These results indicate that an *in vitro* pathological system should not only reproduce a stationary TSS field, but also allow controlled oscillatory motion of the TSS field with relevant frequency and shear magnitude.

**Fig. 2.**
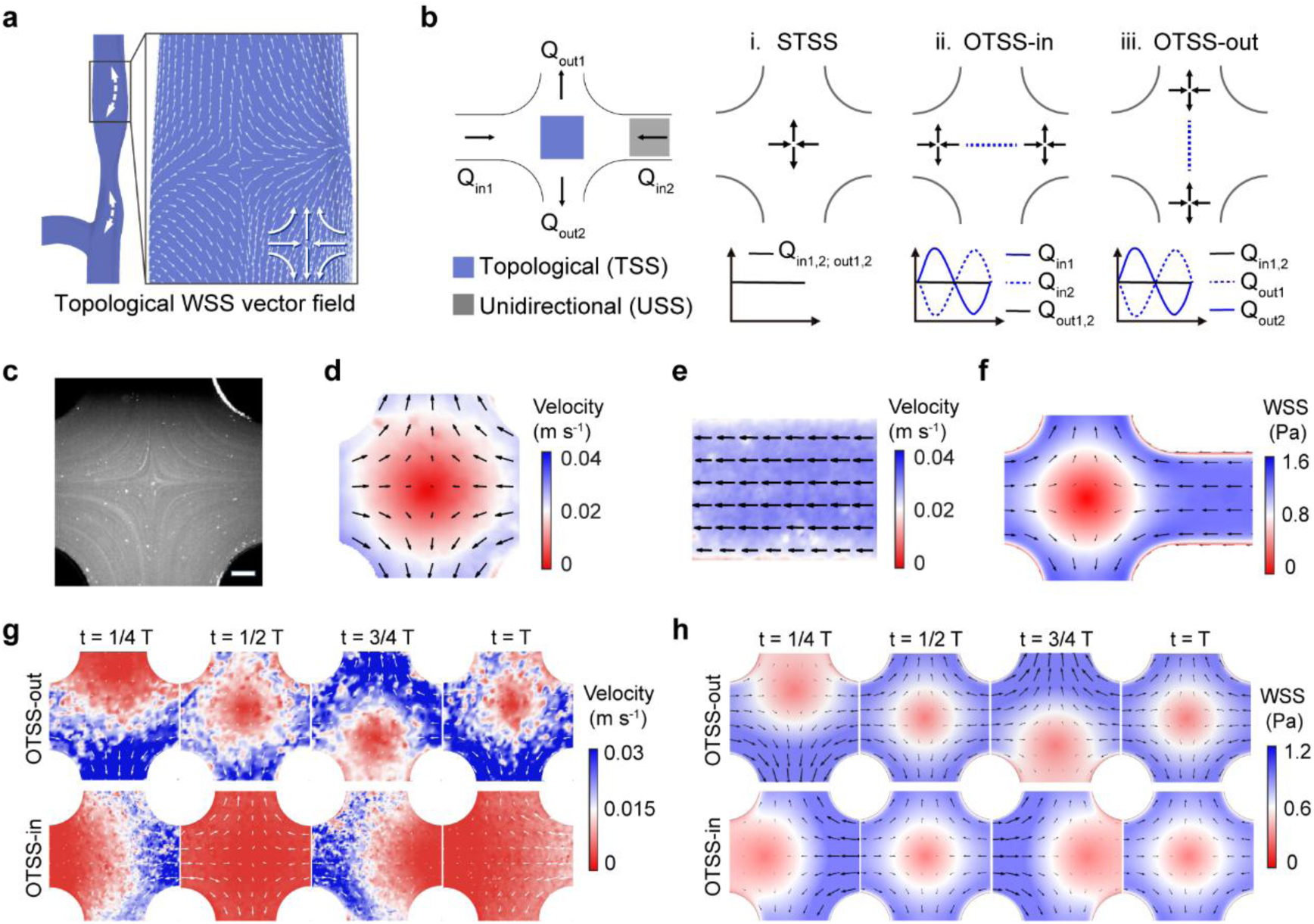
Control and validation of topological flow shear stress in the vascular chip. **a**, Patient-specific simulation of the wall shear stress (WSS) vector field in a stenotic coronary artery. The magnified map shows a topological WSS pattern near the stenotic region. White double-headed arrows indicate upstream and downstream regions where oscillatory displacement of the topological WSS pattern was identified over the cardiac cycle. **b,** Flow configuration of the cross-junction vascular chip. The junction region is exposed to topological shear stress (TSS), whereas the straight-channel regions provide unidirectional shear stress (USS) references. Flow-rate schemes are shown for stationary topological shear stress (STSS), inlet-axis oscillatory topological shear stress (OTSS-in) and outlet-axis oscillatory topological shear stress (OTSS-out). Dashed blue lines indicate the oscillation trajectory of the TSS field. **c,** Representative streamline image in the cross-junction region. Scale bar, 200 μm. **d,e,** μPIV-measured velocity fields in the TSS region under STSS (**d**) and in the USS reference region (**e**). **f,** Simulated WSS magnitude and vector field under STSS, showing the topological shear region at the junction and unidirectional shear in the straight channel. **g,** μPIV-measured velocity fields over one oscillation period for OTSS-out and OTSS-in. **h,** Simulated WSS fields over one oscillation period for OTSS-out and OTSS-in, showing the corresponding direction-specific oscillatory TSS patterns.

To recapitulate these key spatial and temporal features of pathological TSS *in vitro*, we developed a microfluidic vascular chip that reproducibly generates a −1 TSS field at the junction (Fig. 2b, Table S2). By controlling inlet and outlet flow rates, the chip establishes three defined shear conditions: STSS, inlet-axis oscillatory TSS (OTSS-in) and outlet-axis oscillatory TSS (OTSS-out). Under STSS, balanced flow rates maintain the TSS field near the junction center. Under OTSS-in or OTSS-out, sinusoidal modulation of paired inlet or outlet flow rates with a phase difference of π drives the TSS field to oscillate along the corresponding channel axis at 1 Hz, approximating the cardiac cycle. The straight-channel regions maintain unidirectional shear stress (USS) references under matched culture conditions, enabling direct comparison between TSS and USS within the same device. The prescribed flow-rate waveforms were implemented using a computer-controlled perfusion system (Fig. S2a, b)^18^.

The designed flow configurations were then examined experimentally and computationally. Representative streamline imaging visualized converging and diverging flow organization in the cross-junction region (Fig. 2c). Micron-resolution particle image velocimetry (μPIV) measurements resolved the time-averaged velocity field in the central TSS region under STSS and in the straight-channel USS reference region, confirming that the same chip provides both topological and unidirectional shear environments (Fig. 2d, e). Numerical simulations based on the prescribed flow conditions further provided the corresponding WSS magnitude and vector distributions (Fig. 2f). For oscillatory conditions, time-resolved μPIV measurements captured direction-specific displacement of the flow pattern under OTSS-out and OTSS-in over one oscillation period (Fig. 2g). Corresponding numerical analysis of WSS profiles along the channel axis further quantified the displacement range of the TSS field over one oscillation cycle (Fig. 2h, Fig. S2c and Supplementary Video 2). Together, these results establish a controllable vascular-chip system that converts disease-associated WSS topology into defined STSS and OTSS conditions while retaining matched USS references for subsequent endothelial studies.

### Oscillatory topological shear destabilizes collective dynamics of endothelial crowds

Using the validated vascular chip, we next examined endothelial collective dynamics under OTSS, which refers to OTSS-out unless otherwise specified in this section.

Live-cell imaging showed that STSS largely preserved a continuous and relatively uniform endothelial monolayer, whereas OTSS progressively compromised monolayer integrity and induced reduced cell density, intercellular gaps and spatially heterogeneous organization (Fig. 3a and Supplementary Video 3). When quantified as relative cell loss normalized to the initial cell number, OTSS resulted in a markedly greater loss of cells than STSS (Fig. 3b), establishing a structural basis for collective reorganization. To quantify the emergent spatial disorder, we analyzed cell number fluctuations using giant number fluctuation (GNF) analysis^39^. The normalized fluctuation followed a power-law relationship, Δ*N*/√*N* ∝ *N^β^*, where *N* denotes the mean cell number and the scaling exponent *β* quantifies spatial heterogeneity (Fig. 3c and Table S3). Compared with the near-uniform fluctuation profile under STSS, OTSS drove the endothelial layer into a heterogeneous regime with an increased *β*, indicating enhanced spatial disorder^40,41^.

**Fig. 3.**
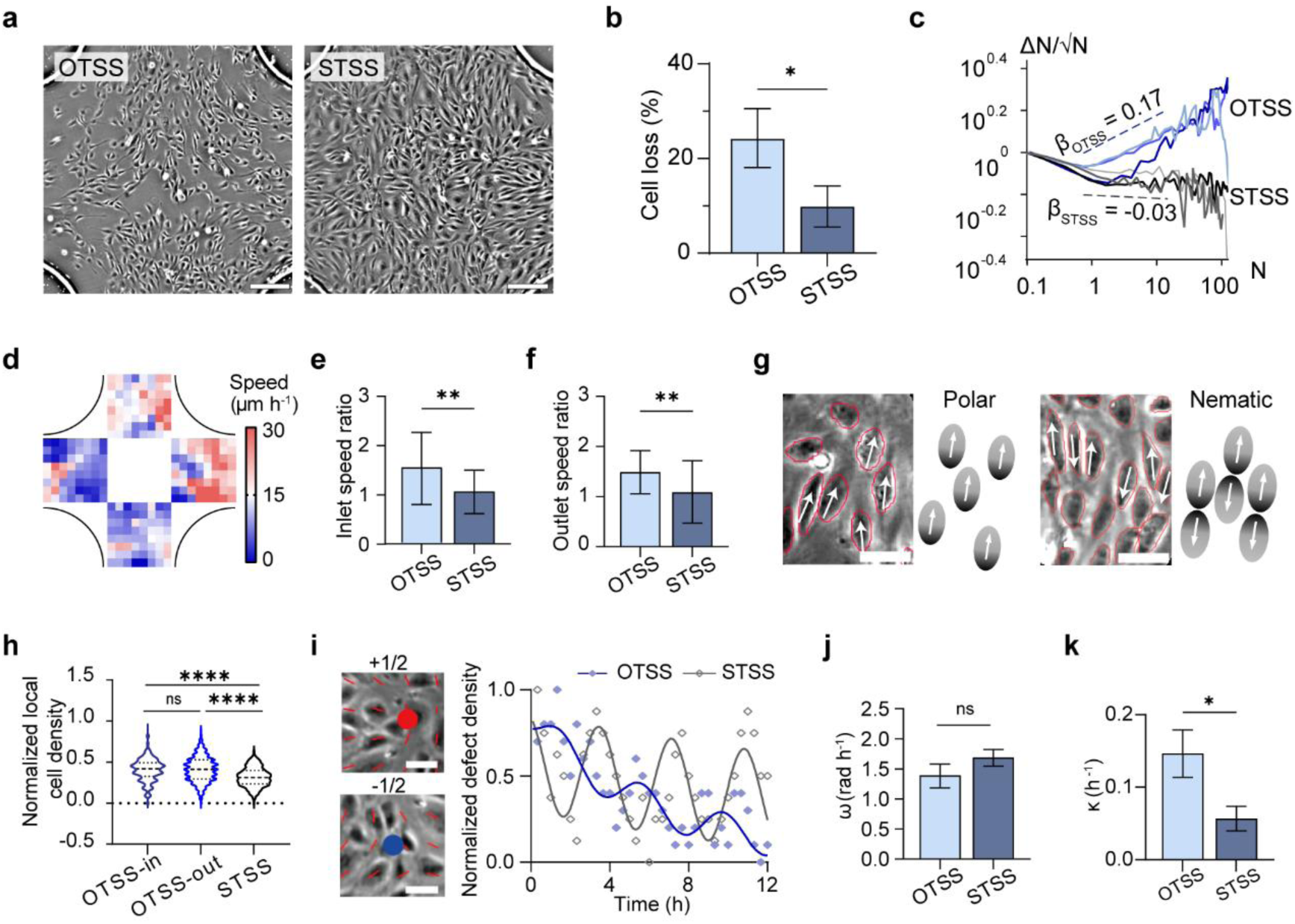
Oscillatory topological shear destabilizes collective dynamics of endothelial crowds. **a**, Representative live-cell images of endothelial monolayers after exposure to oscillatory topological shear stress (OTSS) or stationary topological shear stress (STSS). Scale bars, 200 μm. **b,** Relative cell loss after shear exposure, normalized to the initial cell number. **c,** Giant number fluctuation analysis of endothelial spatial organization under OTSS and STSS. The normalized fluctuation (*ΔN/√N*) is plotted as a function of the mean cell number (*N*), and the fitted scaling exponent *β* is used to quantify spatial heterogeneity. **d,** Representative spatial map of collective migration speed after OTSS exposure. **e,f,** Ratios of mean migration speeds between paired inlet channels (**e**) and paired outlet channels (**f**), showing migration asymmetry under topological shear. **g,** Representative polar and nematic migration patterns identified from live-cell imaging based on local velocity organization. **h,** Normalized local cell density within low-order migration regions under OTSS-in, OTSS-out and STSS. **i,** Representative +1/2 and −1/2 defects in the endothelial orientation field and temporal evolution of normalized defect density under OTSS and STSS. Scale bars, 50 μm. **j,k,** Fitted oscillation frequency *ω* (**j**) and decay coefficient *κ* (**k**) of defect-density dynamics under OTSS and STSS.

This increased heterogeneity was accompanied by asymmetry in collective migration. Despite the geometric symmetry of the chip, OTSS produced inhomogeneous spatial distributions of migration speed across the cross-junction region (Fig. 3d). To quantify this effect, we compared time-averaged migration speeds between geometrically opposing peripheral regions of the cross-junction (Fig. S3a). These ratios report the relative magnitude of collective migration along the inlet and outlet channels (Fig. 3e, f). Under OTSS, both inlet and outlet speed ratios deviated from unity, indicating spontaneous symmetry breaking between opposing sides of the TSS region. In contrast, STSS maintained ratios close to 1, consistent with spatially symmetric migration. Representative single-cell trajectories provided a complementary view of endothelial migration (Fig. S3b, c). Thus, OTSS not only enhanced local heterogeneity but also converted collective migration into a net macroscale asymmetry.

At the mesoscale, OTSS further reorganized endothelial migration into distinct local collective states. OTSS drove endothelial crowds into a phase-separated state characterized by the coexistence of polar and nematic order (Fig. 3g and Supplementary Video 4). This classification was supported by combining a velocity-based order parameter with local cell-density measurements, and by quantifying local density in low-order migration regions (Fig. 3h). In cell-sparse regions, endothelial collectives adopted a polar state, characterized by strong local velocity alignment and coherent unidirectional migration. In contrast, denser regions self-organized into a nematic state, in which reduced velocity alignment was accompanied by persistent antiparallel streams, resulting in orientational alignment but near-cancellation of the coarse-grained migration polarity. Time-lapse imaging further captured local antiparallel rearrangement and cell extrusion under OTSS, illustrating a mechanical basis for density loss in nematic regions as neighboring cells reorganized (Fig. S3d). Such arrangements are consistent with the antiparallel cell streams under confinement and angiogenic morphogenesis^31,42,43^. These observations indicate that OTSS selects local active phases through the coupled effects of TSS and cell density. This density–order coupling provides a quantitative basis to distinguish polar versus nematic collective states in a population of intrinsically polar, self-propelled cells.

We next analyzed the dynamics of ±1/2 topological defects in the endothelial orientation field. Such half-integer defects, defined by characteristic winding numbers in the cell orientation field, provide a metric for collective reorganization in ordered cell layers. Representative +1/2 and −1/2 defects were identified from the orientation field, and their normalized density was tracked over time (Fig. 3i). The temporal evolution of normalized defect density was fitted to a damped oscillatory function,

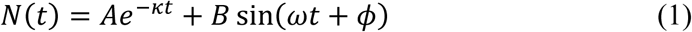

where *κ* is the decay coefficient and *ω* is the frequency of the oscillatory component (Table S4). Although the fitted oscillation frequency remained comparable between OTSS and STSS (Fig. 3j), the decay coefficient was significantly larger under OTSS (Fig. 3k). This suggests that OTSS preserves a comparable oscillatory rhythm but shortens defect lifetime and accelerates the cycle of creation and annihilation, leading to repeated collective reorganization rather than monotonic relaxation^44,45^.

Moreover, we examined whether the direction of TSS oscillation further modulated endothelial collective dynamics. OTSS-in induced cell detachment and spatial heterogeneity comparable to OTSS-out (Fig. S4a–d). However, extended migration analyses showed that OTSS-in reduced average migration speed and persistence compared with OTSS-out (Fig. S3a and Table S5), and defect-density fitting revealed direction-dependent changes in oscillation frequency and decay coefficient (Fig. S4e, f and Table S4). Together, these results show that temporal oscillation and spatial directionality of topological shear jointly destabilize and reorganize endothelial collective dynamics.

### Oscillatory topological shear impairs actin–nuclear mechanotransduction

Having established the collective consequences of OTSS, we next examined how these physical perturbations translate into altered mechanobiological function^11^. To this end, we investigated the actin–nuclear axis, a primary mechanotransduction linkage where the perinuclear actin architecture couples with the nucleus to maintain stability^32,34^. We focused on the perinuclear actin cap, which is positioned apically above the nucleus and contributes to nuclear shaping and mechanical shielding.

Because nuclear mechanotransduction is also coupled to transcriptional mechanoregulators, we further examined YAP localization as an indicator of mechanotransduction-associated nuclear signaling. Accordingly, we analyzed F-actin organization, nuclear morphology, chromatin condensation, nuclear mechanics and YAP distribution in endothelial cells exposed to defined shear conditions (Fig. 4a).

**Fig. 4.**
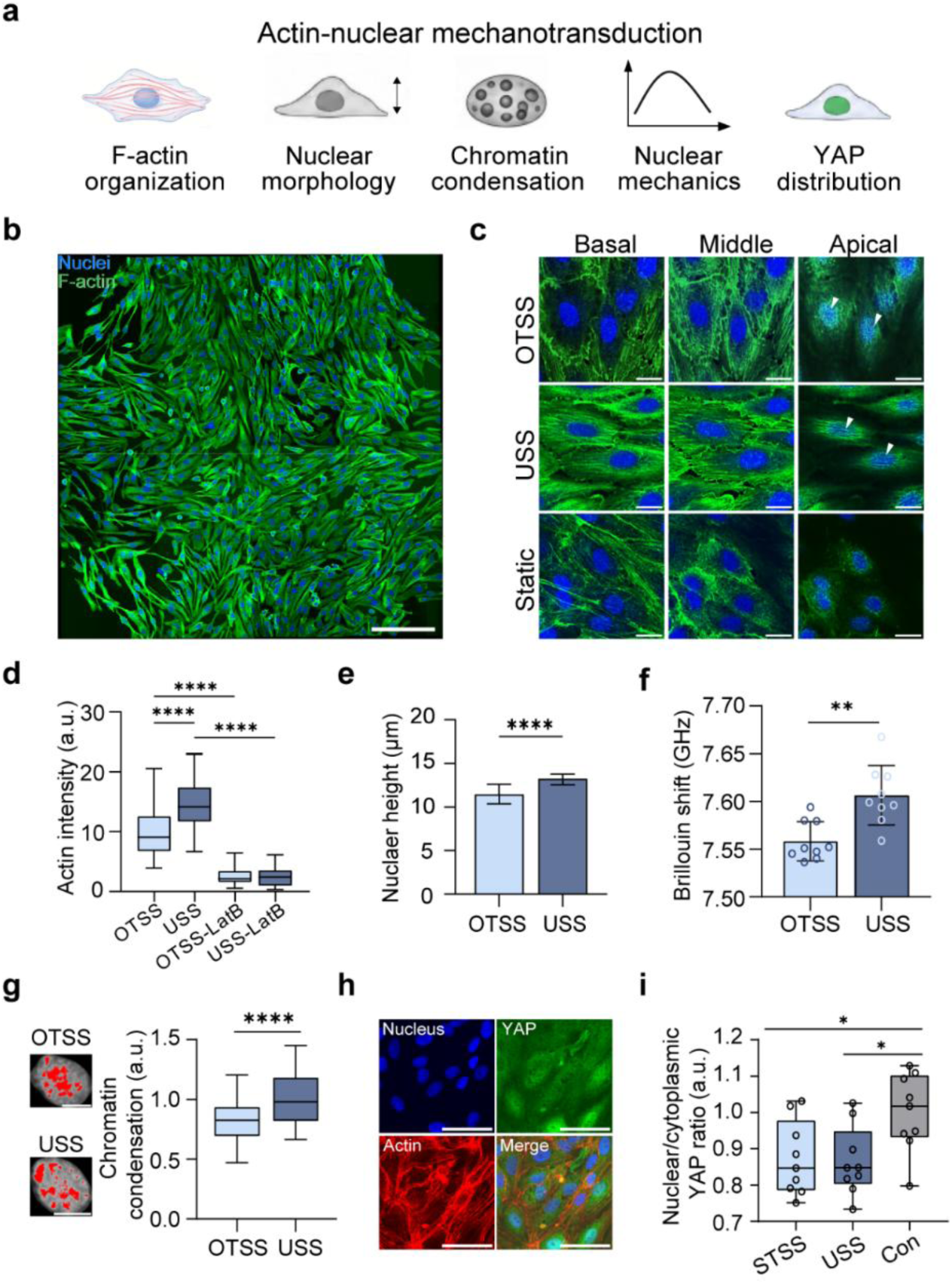
Oscillatory topological shear impairs actin–nuclear mechanotransduction. **a**, Schematic overview of actin–nuclear mechanotransduction and the mechanobiological parameters examined in this study, including F-actin organization, nuclear morphology, chromatin condensation, nuclear mechanics and YAP distribution. **b**, Stitched fluorescence image of endothelial cells exposed to oscillatory topological shear stress (OTSS), showing nuclei and F-actin. Scale bar, 200 μm. **c**, Representative images of endothelial cells under OTSS, USS and static conditions at basal, middle and apical planes, showing nuclei and F-actin. White arrowheads indicate apical actin-cap structures. Scale bars, 20 μm. **d**, Quantification of actin intensity under OTSS, USS and Latrunculin B-treated conditions. **e**, Quantification of nuclear height under OTSS and USS. **f**, Nuclear Brillouin frequency shift under OTSS and USS, used as a label-free indicator of nuclear mechanical properties. **g**, Representative chromatin-condensation maps and quantification of chromatin condensation under OTSS and USS. **h**, Representative fluorescence images showing nuclei, YAP, F-actin and merged channels. Scale bars, 50 μm. **i**, Quantification of the nuclear/cytoplasmic YAP ratio under stationary topological shear stress (STSS), USS and static control conditions.

Stitched fluorescence imaging first showed the spatial organization of nuclei and F-actin in endothelial monolayers exposed to OTSS (Fig. 4b). Layer-resolved fluorescence imaging across basal, middle and apical planes further revealed distinct actin organization under different shear conditions (Fig. 4c). Cells under USS formed robust apical actin caps above the nucleus, whereas cells under OTSS exhibited disrupted and less organized actin fibers. Z-resolved analysis of the apical tier was used to quantify actin-cap integrity, with the analysis region defined using the nuclear mask (Fig. S5a). Quantitative analysis confirmed reduced actin intensity under OTSS and Latrunculin B (LatB) treated conditions compared with USS (Fig. 4d). Consistent with impaired cytoskeletal support, cells under OTSS exhibited pronounced nuclear flattening, evidenced by a marked reduction in nuclear height relative to USS (Fig. 4e).

We next examined whether these cytoskeletal and nuclear-shape changes were accompanied by altered nuclear mechanical properties. Non-invasive Brillouin microscopy showed that cells exposed to OTSS exhibited a reduced nuclear Brillouin frequency shift relative to USS, indicating a lower apparent nuclear mechanical modulus and a softer nucleus (Fig. 4f, Fig. S5b-d). Chromatin-condensation analysis further showed reduced chromatin condensation under OTSS compared with USS, suggesting a less compact nuclear state (Fig. 4g)^46,47^. Finally, fluorescence imaging and quantification of YAP localization showed that both STSS and USS reduced the nuclear/cytoplasmic YAP ratio relative to static control conditions, whereas no marked difference was observed between STSS and USS (Fig. 4h, i). Together, these results indicate that OTSS disrupts actin-cap organization and is associated with nuclear flattening, reduced nuclear mechanical properties and decreased chromatin condensation, while the chip also enables analysis of mechanotransduction-associated YAP distribution under defined shear-topology conditions.

### Oscillatory topological shear promotes endothelial nanoparticle uptake

We next asked whether the structural and mechanical changes induced by OTSS were accompanied by altered endothelial transport-related function^12^. Because the endothelium acts as a key gateway for vascular drug delivery, nanoparticle (NP) uptake provides a functional assay of endothelial transport permissiveness. Endothelial monolayers were first exposed to defined TSS conditions, then incubated with fluorescent NPs, and NP accumulation was quantified using a perinuclear region (Fig. 5a, Fig. S5a).

**Fig. 5.**
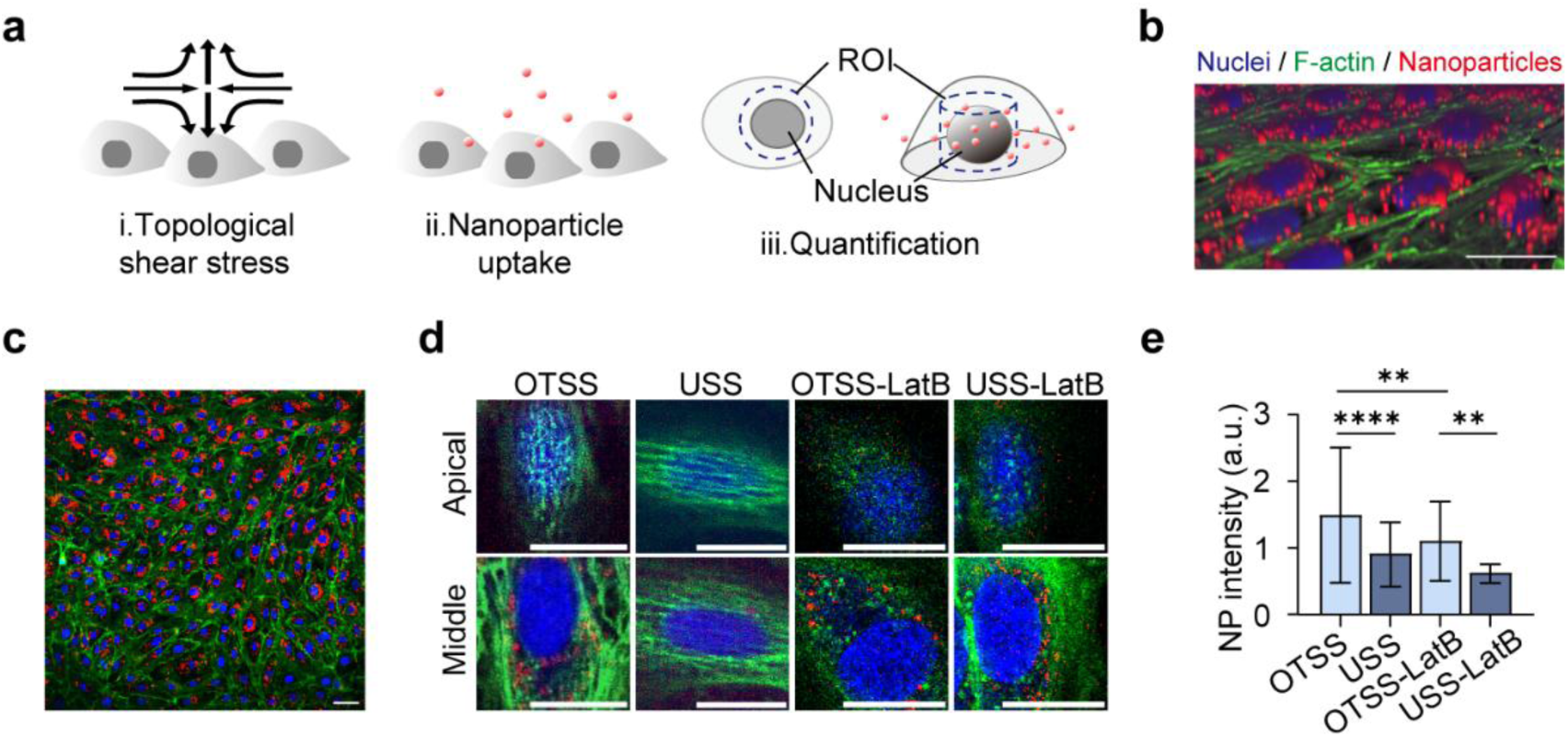
Oscillatory topological shear promotes endothelial nanoparticle uptake. **a**, Schematic overview of the nanoparticle-uptake experiment, including i, exposure to topological shear stress; ii, incubation with fluorescent nanoparticles and endothelial uptake; and iii, quantification of nanoparticle accumulation using a perinuclear region of interest (ROI). b, Three-dimensional reconstructed fluorescence image showing nanoparticle distribution relative to endothelial nuclei and F-actin. Nuclei are shown in blue, F-actin in green and nanoparticles in red. Scale bar, 50 μm. c, Stitched fluorescence image showing endothelial nanoparticle uptake under oscillatory topological shear stress (OTSS). Scale bar, 50 μm. d, Representative fluorescence images at the apical and middle planes under OTSS, unidirectional shear stress (USS), OTSS with Latrunculin B treatment (OTSS-LatB) and USS with Latrunculin B treatment (USS-LatB). Scale bars, 20 μm. e, Quantification of perinuclear nanoparticle fluorescence intensity under OTSS, USS, OTSS-LatB and USS-LatB conditions.

Three-dimensional fluorescence reconstruction showed NP distribution relative to endothelial nuclei and F-actin, revealing that fluorescent polystyrene NPs were internalized and preferentially accumulated in the perinuclear region (Fig. 5b).

Stitched fluorescence imaging further showed NP uptake across the OTSS-exposed endothelial monolayer (Fig. 5c). Layer-resolved fluorescence imaging at apical and middle planes revealed stronger NP signals under OTSS than under USS, with NPs preferentially accumulating near the perinuclear region (Fig. 5d). Quantitative analysis confirmed increased perinuclear NP fluorescence intensity under OTSS compared with USS (Fig. 5e), indicating that OTSS promotes a transport-permissive endothelial state.

To test whether actin dynamics contributed to this enhanced uptake, we perturbed filament assembly using LatB. Under OTSS, LatB treatment attenuated the elevated perinuclear NP accumulation, whereas its effect under USS remained limited (Fig. 5d, e). These results indicate that OTSS-enhanced NP accumulation involves an active, actin-dependent process, consistent with established models in which actin polymerization facilitates endocytic internalization by counteracting membrane tension^48^. Together with the actin–nuclear disruption observed above, these findings link oscillatory topological shear to altered endothelial nanoparticle uptake.

## Discussion

Our study establishes a vascular-chip strategy in which disease-relevant WSS topology functions as an independently programmable experimental parameter. Through spatiotemporal recapitulation of the −1 TSS, we transform a hemodynamic feature previously accessible mainly through computational inference into a controllable input that can be stabilized as stationary TSS (STSS), dynamically modulated into oscillatory TSS (OTSS), and directly benchmarked against matched unidirectional shear stress (USS) within a single standardized device. This enables systematic interrogation of how a single pathological mechanical signature propagates across endothelial collective organization, subcellular mechanotransduction, and transport function, providing a unified framework for cross-scale vascular phenotyping.

The present vascular-chip strategy addresses a limitation shared by both hemodynamic descriptors and experimental vascular models. This two-dimensional topological flow architecture is fundamentally distinct from conventional mechanobiological platforms, including uniform shear chambers, step-flow systems, and sequential multi-directional flow devices, which constrain shear organization through fixed geometries^6,49,50^. At the same time, the system remains compatible with standard organ-on-chip platforms, which has potential implications for drug screening and mechanomedicine^51,52^. Importantly, this design allows WSS topology to be independently modulated without altering local shear magnitude, enabling direct isolation of topology as an experimental variable.

Precise control over WSS topology also reframes the mechanical interpretation of endothelial collective dysfunction. By capturing real-time cellular responses to the complex flow topology, our results demonstrate that OTSS alone is sufficient to destabilize monolayer integrity and drive asymmetric collective dynamics under matched shear magnitude, indicating that endothelial responses depend not only on shear intensity, but also on the spatial organization and temporal evolution of the shear-vector field. We further identify cell density as a critical modulator of collective dynamics. Under OTSS, cell density is not constant but evolves dynamically and is crucial for the observed collective behavior. At high cell density, the intrinsic polarity reversal and mutual collisions of elongated cells drive local nematic alignment, leading to a phase landscape comprising polar, nematic, and coexistence states^53^. This is consistent with recent findings that non-monotonic evolution of nematic topological defects of endothelial layer was only observed at high cell densities and when the crowd density reaches a threshold^30^, groups of confined people can undergo self-sustained oscillations^54^. The methodology can further be extended to explore coupling between WSS topology and cellular collective behaviors, a correspondence that has remained largely inaccessible in conventional systems.

At the subcellular scale, OTSS propagates through the actin–nuclear mechanotransduction axis. It disrupts perinuclear actin organization, alters nuclear morphology, and reduces nuclear stiffness, consistent with established roles of actin remodeling, perinuclear actin structures, and nuclear mechanics in shear-induced mechanotransduction^55,56^. Since WSS can simultaneously regulate cellular activity and alignment through mechanobiological pathways such as the actin–nuclear axis, the present study couples intrinsic cellular activity with flow-induced alignment, rather than modeling these effects separately as in previous studies^30,31^. The integration of collective cell migration, fluorescence-based structural readouts with label-free Brillouin microscopy enables quantitative coupling between mechanical input and nuclear mechanical state. Such cross-scale integration is central to mechanomedicine, where mechanical signatures are increasingly viewed as measurable phenotypes, causal regulators, and therapeutic targets^26–28,52^.

The nanoparticle uptake results extend this framework from mechanistic vascular biology toward functional screening. Vascular-targeted delivery depends on near-wall transport, endothelial barrier state, cytoskeletal organization and intracellular uptake kinetics^35,48,57^. Previous work under conventional unidirectional shear configurations has shown that elevated flow rates reduce endothelial nanoparticle uptake^33^. In contrast, OTSS enhances perinuclear nanoparticle accumulation relative to matched USS, indicating vascular transport permissiveness cannot be inferred from shear magnitude or flow rate alone, and that WSS spatial organization generates topology-dependent endothelial states that alter particle access and intracellular distribution. Programmable topological shear thus provides a disease-relevant test condition for evaluating nanotherapeutics and vascular-targeted agents.

The engineered −1 TSS field represents a reduced, standardized instantiation of a recurrent pathological motif, rather than an anatomically exhaustive disease model. Future work will extend this strategy to additional WSS topologies found at vascular bifurcations, stenoses, aneurysms and stented segments, and incorporate vascular smooth muscle cells, immune cell interactions, extracellular matrix remodeling, inflammatory cues and vessel-wall compliance. Because the platform decouples topology from anatomy, these extensions can be implemented within a standardized readout pipeline rather than requiring bespoke disease-specific models. Ultimately, this work underscores the necessity of recapitulating physiologically and pathologically relevant mechanical environments, beyond idealized uniform flow, to uncover genuine mechanisms of tissue organization. It provides an experimental framework for translating computational hemodynamic topology into standardized vascular disease modeling, mechanomedicine-oriented analysis and topology-informed therapeutic screening.

## Methods

### Fabrication of microfluidic chip

Microfluidic molds were fabricated using standard soft lithography. Devices were cast in polydimethylsiloxane (PDMS; Sylgard 184, Dow Corning) with a 10:1 ratio of PDMS base to curing agent, and irreversibly bonded to glass coverslips using oxygen plasma treatment. The device featured a cross-junction that connected two inlets and two outlets, enabling programmable generation of wall shear stress (WSS) vector fields with a −1 defect (saddle fixed point) topology. The oscillatory topological shear stress (OTSS) region was confined to the cross-junction, while the straight inlet and outlet channels acted as control regions subjected to steady unidirectional shear stress (USS) at physiological magnitude 1.0 Pa. The inlet and outlet channels had a width of 800 μm, and the cross-junction area spanned an approximate 1000 × 1000 μm field of view. The overall chip height was 120 μm. Three flow conditions were implemented: stationary topological shear stress (STSS) with a fixed defect at the junction center, OTSS-in with defect oscillation along the inlet axis, and OTSS-out with defect oscillation along the outlet axis (Fig. 2b). Unless specified, OTSS refers to OTSS-out.

### Automated flow control and PIV validation

Flow fields were generated by coordinating multiple syringe pumps using a custom automation program that executed predefined flow-rate waveforms. Under OTSS conditions, opposing waveforms were sinusoidally modulated with a phase shift of π at 1 Hz, while maintaining a constant mean flow rate to achieve physiologically relevant WSS levels in the straight channels (Supplementary Table S2). Additional implementation details of the automated perfusion system are available in our previously published study^18^.

To validate the flow topology, micron-resolution particle image velocimetry (μPIV; microVec TRSM-1M9000-20W, China) was performed. Fluorescent tracer particles with 2 μm diameter were suspended in culture medium. Time-resolved images were acquired at 2000 fps under the same waveforms as in cell experiments. Velocity fields were computed in PIVlab (v2.55) running on MATLAB (R2021a), using a 32 × 32 pixel interrogation window^58^. For each condition, more than 100 image pairs were analyzed. Near-wall velocity gradients were used to infer WSS patterns and verify the expected unidirectional and topological shear stress distributions (Fig. 2d, e and g).

### Cell culture and seeding

Human umbilical vein endothelial cells (HUVECs) were isolated from newborn umbilical cords as described previously^16^. Informed consent was obtained from all donors, and procedures were approved by the Ethics Committee of Beihang University. Primary HUVECs were cultured in endothelial medium consisting of a 1:1 mixture of M199 (Gibco, Cat. 31100035) and ECM (ScienCell, Cat. 43001) supplemented with 10% fetal bovine serum. Microfluidic chips were sterilized by UV irradiation for 1 h, coated with fibronectin (80 μg mL⁻¹; Corning, Cat. 356008) at 37 °C for 2 h, and seeded with HUVECs (passages 3–6). Monolayers were allowed to reach confluence for about 5 hours and then subjected to fluid flow for 12 hours in a stage-top incubator maintained at 37 °C and 5% CO₂. For actin perturbation experiments, Latrunculin B (LatB, 60 μM) was introduced during shear exposure.

### Nanoparticle uptake assay

Following shear exposure, fluorescent polystyrene nanoparticles (200 nm diameter, Invitrogen, Cat. 2451250) were diluted 1:5000 in culture medium and slowly injected into the microfluidic channels to ensure complete filling^59^. Chips were incubated at 37 °C for 4 hours to allow nanoparticle uptake. Samples were then fixed with 4% paraformaldehyde and processed for fluorescence imaging.

### Live-cell imaging

Collective endothelial dynamics were recorded by phase-contrast time-lapse microscopy (Leica DMi8, Germany) equipped with a 10× objective and a Leica DFC9000 GT camera. Imaging was focused on the cross-junction region, with a single field of view spanning the full channel width. Images were acquired every 20 minutes from the onset of shear exposure to the end of the experiment.

### Image processing and cell tracking

Image sequences were preprocessed in ImageJ through image registration and rotation to align the channel axis. A square region of interest centered on the cross-junction (1000 μm × 1000 μm; 800 × 800 pixels) was cropped from each frame. Images were bandpass-filtered with a characteristic scale of 40 pixels and contrast-normalized to enhance cell boundaries. Cell segmentation and tracking were performed in TrackMate (v7.14) ^60^, integrated with Cellpose (v2.0) for cell segmentation^61^. For tracking, the cytoplasm model and grayscale channel were used, with an expected cell diameter of 30 pixel. Ellipse-fit parameters exported from TrackMate were used to quantify cell count, area, aspect ratio, and the long-axis orientation of each segmented cell. Cell migration speed was computed from centroid displacements between consecutive frames (time interval = 20 minutes). Migration speed was quantified within spatial subregions defined by a 3 × 3 partition of the 1000 μm field of view. Each tile measured 300 μm × 300 μm. The four side-adjacent tiles corresponding to the two inlet and two outlet straight channels were used to calculate area-averaged migration velocities for each channel segment, denoted as *u_in_*_1_, *u_in_*_2_, *u_out_*_1_, and *u_out_*_2_, respectively. The central tile containing the −1 TSS core was used to compute the area-averaged central velocity, denoted as *u_cen_*. Stable migration speed was defined as the time-averaged speed during the final one-third of flow stimulation. The mean peripheral velocity was defined as the average of the four peripheral region migration speeds (*u_in_*_1_, *u_in_*_2_, *u_out_*_1_, and *u_out_*_2_). Speed ratios were computed either between paired inlet or outlet regions or between the mean peripheral velocity and the central speed *u_cen_*, as indicated in Fig. S3a. For ratio-based analyses, values were normalized such that ratios ≥ 1 were retained (i.e., the larger regional velocity was divided by the smaller), enabling symmetric comparison of bilateral or peripheral–central differences. Deviation of the ratio from 1 was interpreted as spatial heterogeneity in migration speed.

### Cell Displacement Analysis

To analyze cell migration, we utilized a model based on persistent random walk behavior, which is commonly used to describe collective motion in biological systems (Supplementary Table S5)^62^. Mean squared displacement of cells at time *t* was fitted using the following equation:

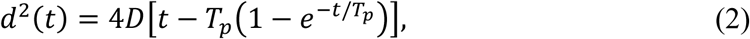

where *D* is the diffusion coefficient that characterizes the long-term random motion of the cell. It indicates the rate at which the cell’s displacement spreads over time, representing the diffusivity of the cell’s movement over extended periods. *T_p_* is the persistence time, representing the characteristic time for which the cell maintains its directionality during movement. A higher *T_p_* indicates that the cell moves in a more directional manner, while a smaller *T_p_* corresponds to a more random movement.

### Velocity-Based Order Parameter

The velocity-based order parameter quantifies the local collective organization of cells by comparing the velocity directions of a given cell with its surrounding neighbors within a specified radius. The order parameter *S* for a given cell *i* is calculated as:

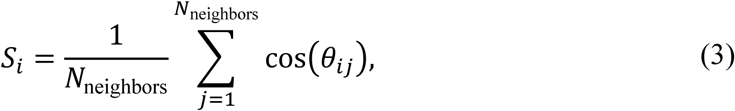

where *θ_ij_* is the angle between the velocity vector of cell *i* and the velocity vector of its neighbor *j*, and *N*_neighbors_ is the number of neighboring cells within a defined radius (typically 50 μm). The cosine of the angle is used to measure the similarity of the velocity direction. The same neighborhoods were used to estimate local cell density by counting segmented cells within the radius.

### Giant number fluctuation analysis

Spatial heterogeneity was quantified by giant number fluctuations (GNF) computed from cell counts within square sampling windows placed across the analysis region. The region of interest was partitioned into square windows spanning side lengths from 5 to 500 pixels. For each window size, the mean cell count *N* and standard deviation Δ*N* were computed, and the normalized fluctuation metric was fitted to a power law:

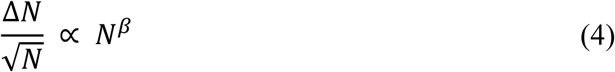

on log–log axes to obtain the scaling exponent *β*, following established GNF analyses in active matter and cell collectives^39^. The fitting range for *N* (1–10) and regression procedure are detailed in Supplementary Table S3.

### Orientation field construction and topological defect identification

Topological defects were identified from cell orientation fields following established methods^63^. Orientation fields were computed from phase-contrast images using OrientationJ with a grid size of 12 pixels^64^. The local nematic Q-tensor was computed and defect were classified by evaluating the winding number. Defect density was defined as the number of detected ±1/2 defects within the region of interest divided by the analyzed area. Temporal defect density *N*(*t*) was computed at each time point and normalized by the initial value for each experiment. The time evolution of normalized defect density was fitted to a damped oscillatory function,

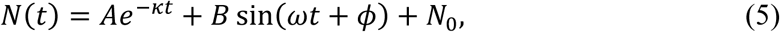

where *κ* is the decay coefficient, *ω* is the angular frequency, *A* and *B* are fitted scaling coefficients, and *N*_0_ is the steady-state offset. Fit parameters were obtained by nonlinear least squares regression, and fits with R^2^ ≥ 0.6 were retained for analysis (Supplementary Table S4).

### Brillouin microscopy

Nuclear mechanical properties were measured in live cells immediately after shear exposure using a custom-built Brillouin microscope as previously described^34^. A 532 nm continuous laser was focused through a 60× water-immersion objective (UPlanSApo 60×W, NA 1.2; Olympus, Japan) and Brillouin spectra were analyzed by a VIPA-based spectrometer. A bright-field imaging channel was used to locate nuclei prior to Brillouin acquisition. Brillouin signals were acquired using line scans across individual cells with 100 measurement points at 0.5 μm spacing (50 μm total length). Brillouin spectra were processed using custom MATLAB software that synchronized stage/camera control and performed automated least-squares Lorentzian fitting. Fits with *R*^2^ ≥ 0.8 were retained. Nuclear Brillouin shifts were computed as the mean of at least 15 points within the nuclear region. For each flow condition, 3 independent samples were measured with more than 3 cells per sample (Fig. 5b-d).

### Fluorescence staining and confocal microscopy

Cells were fixed in 4% paraformaldehyde for 10 minutes, permeabilized with 0.1% Triton X-100 in PBS for 15 minutes, and blocked with 5% bovine serum albumin for 1 hour at room temperature. F-actin was stained with YF488-conjugated phalloidin (1:200; UElandy, Cat. YP00595), and nuclei were counterstained with DAPI (1:500; Sigma–Aldrich) for 10 minutes. All fluorescence imaging was performed using a spinning-disk confocal microscopy system (Andor Dragonfly mounted on a Leica platform) equipped with a 40× oil-immersion objective. Confocal z-stacks were acquired across the full cell height with a z-step of 0.3 μm. For layer-resolved representations, the fluorescence-defined cell height (bottom to top) was divided into basal, middle, and apical thirds, and the central slice of each third was used as the representative basal/middle/apical plane. Apical stacks were used for actin cap imaging (Fig. 5a). Chromatin compaction was quantified by the coefficient of variation (CV) of DAPI intensity within each segmented nucleus, following established microscopy-based methods. Actin cap intensity was quantified from the apical third of confocal z-stacks by generating an intensity projection of the phalloidin channel restricted to apical planes above the nucleus to minimize basal fiber contributions. The mean F-actin intensity within the corresponding nuclear footprint was computed for each cell. Nanoparticle (NP) uptake was quantified from the middle focal plane by measuring NP channel fluorescence intensity within a perinuclear region defined from the nuclear mask after two erosion operations. To reduce background contributions, the mean fluorescence intensity of cell-free regions in each image was used as the background level and subtracted from per-cell measurements. Nanoparticle uptake was reported as fluorescence intensity normalized by the analyzed area to reduce cell-size dependence. For each flow condition, at least three independent samples were analyzed, with more than 30 cells quantified per sample.

### Numerical simulation of a stenotic left coronary artery

A patient-specific stenotic left coronary artery model was constructed from digital subtraction angiography (DSA) image sequences, which provided both vessel lumen dimensions and time-resolved contrast propagation for flow waveform extraction. Angiographic images were processed in MATLAB (R2021b) using Laplacian filtering (3×3 kernel), followed by Hessian-based vessel enhancement and region-growing segmentation to extract the inlet flow boundaries. Key vascular dimensions, including the left main artery (LM), left anterior descending artery (LAD), and left circumflex artery (LCX), were extracted and used to construct a three-dimensional bifurcation geometry in SolidWorks (v2021). The diameters of LM, LAD, and LCX are 3.6 mm, 3.5 mm, and 3.0 mm, respectively (Supplementary Table S1). The proximal segment of LM and the distal segments of LAD and LCX were extended by 40 mm to minimize boundary effects. A focal stenosis with a diameter of approximately 2.5 mm was introduced in the proximal LAD. The proximal LM was assigned as the velocity inlet, and the distal LAD and LCX as pressure outlets (Fig. S1a).

Transient blood flow was simulated in COMSOL Multiphysics (v6.2) under the assumption of incompressible laminar Newtonian behavior. The governing equations were the transient Navier–Stokes and continuity equations:

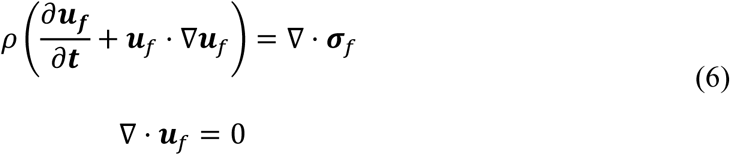

where ***u****_f_* represents the fluid velocity vector, *ρ* =1060 kg·m⁻³ is the fluid density, and ***σ****_f_* represents the fluid stress tensor:

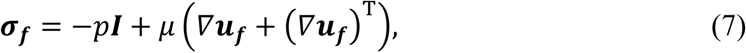

with *p* denoting pressure and dynamic viscosity *μ* = 3.5 × 10^−3^ Pa · s . Vessel walls were modeled as rigid with no-slip boundary conditions^65^. The inlet velocity waveform was extracted directly from the DSA angiographic sequence by tracking contrast-agent propagation along the vessel centerline over one cardiac cycle. The discrete velocity data were smoothed using a fourth-order Fourier fit to generate a continuous periodic inlet waveform. The computational domain was discretized using an unstructured tetrahedral mesh comprising 184,597 vertices and 771,990 elements. Mesh independence was verified by less than 1% variation in time-averaged wall shear stress (WSS) upon refinement. Simulations were performed with a time step of 0.002 s, resolving one cardiac cycle with 394 temporal sampling points. A total of six cardiac cycles were simulated, and results from the final cycle were used for analysis.

To characterize the topological features of the flow, the WSS vector field on the luminal surface was extracted and analyzed using topological skeleton analysis^21^. Fixed points of the WSS vector field were identified and classified based on the eigenvalues of the Jacobian matrix at the fixed-point location. Specifically, fixed points were categorized into three types: saddle points, unstable nodes, and stable focus (Supplementary Video 1). We specifically tracked the trajectory of the downstream saddle point throughout the cardiac cycle to quantify the spatiotemporal range of the oscillatory topological shear stress (Fig. S1b, c).

### Numerical Simulation of Topological Shear Stress Field in Vascular Microfluidic Chips

The three-dimensional flow simulation was conducted using COMSOL Multiphysics (v6.2). Laminar-flow simulations were then performed to parameterize the flow-rate waveforms needed to generate stationary topological shear stress (STSS) and oscillatory topological shear stress (OTSS) on the endothelial monolayer. The microfluidic geometry, consisting of a cross-shaped channel with two inlets and two outlets, was specifically designed to apply different shear stress conditions. This design is shown in Fig. 2b. For STSS, a steady inlet flow rate was applied, with both outlet pressures set to zero. The flow rate was calibrated to achieve a WSS of approximately 1.0 Pa in the straight channels, which is a physiological target for arteries. For OTSS, two distinct configurations were modeled: OTSS-in and OTSS-out. In the OTSS-in configuration, the flow rate of two inlets was sinusoidally modulated at a frequency of 1 Hz with a phase shift of π, while both outlets were held at zero pressure. This setup generated a moving −1 topological defect core that oscillated along the inlet axis. In the OTSS-out configuration, the inlet flow was held constant, and the outlets were modulated sinusoidally with a π phase shift, resulting in an oscillating −1 defect core that moved along the outlet axis. The amplitude of the sinusoidal flow waveforms was adjusted to ensure that the −1 defect core traversed the main cross-junction area during each oscillation cycle. The experimentally used waveform parameters are summarized in Supplementary Table S2. Additionally, fixed point analysis was performed on the flow field using the same methodology applied to the coronary artery stenosis model. Only one saddle point (−1 topological defect) was identified, which was observed to oscillate throughout the oscillation cycle (Supplementary Video 2).

## Statistics and reproducibility

All experiments were performed in at least three independent biological replicates unless stated otherwise. Data are reported as mean ± s.d. Two-sided unpaired Student’s t-tests were used for two-group comparisons. For comparisons among three or more groups, one-way or two-way ANOVA with Tukey’s post hoc test was applied. Normality and variance assumptions were evaluated using Brown–Forsythe or Bartlett tests where appropriate. Statistical significance was defined as *p* < 0.05. No statistical methods were used to predetermine sample size, and investigators were not blinded to experimental conditions.

## Data, code, and materials availability

All data supporting the findings of this study are available from the corresponding authors on request.

## Acknowledgements

We thank all members of the laboratory for their helpful discussions and technical assistance throughout this project. This work was supported by the Beijing Natural Science Foundation (Z260018, L241032 and L246053), the National Key Research and Development Program of China (2025YFC2422603), the National Natural Science Foundation of China (32371375, 31971244, and 12332019), the Fundamental Research Funds for the General Universities, and the Non-profit Central Research Institute Fund of Chinese Academy of Medical Sciences (2024-JKCS-21) .

## Author contributions

X.L. and K.L. conceptualized the study and coordinated the work. K.L., S.Y., K.H., and L.W. performed the investigations and developed the methodology. K.L., K.H., Z.L., and X.Z. conducted the experiments and collected the data. S.Y., U.M., V.M., and D.G. carried out the formal analysis. J.Y., M.W., X.S., Y.Z. and Y.C. provided essential resources and materials. Z.C., A.S., and L.C. provided technical support and assisted with data curation. The original draft of the manuscript was written by K.L. and S.Y., and subsequently reviewed and edited by U.M. and X.L. X.L. supervised the project and acquired the funding. All authors reviewed the manuscript before submission and approved the final version.

## Competing interests

Authors declare that they have no competing interests.

## Additional information

Supplementary information includes Supplementary Figures, Supplementary Tables, and video titles and captions.

Correspondence and requests for materials should be addressed to Kexin Li or Xiao Liu.

## Notes

### Competing Interest Statement

The authors have declared no competing interest.

