## Supplementary Information for "A vascular chip for disease-relevant flow shear stress topology"

**This file includes:**

Supplementary Fig. S1 to Fig. S5
Supplementary Table S1 to S5
Supplementary Video captions including Supplementary Video 1 to 4

**Supplementary Figure**

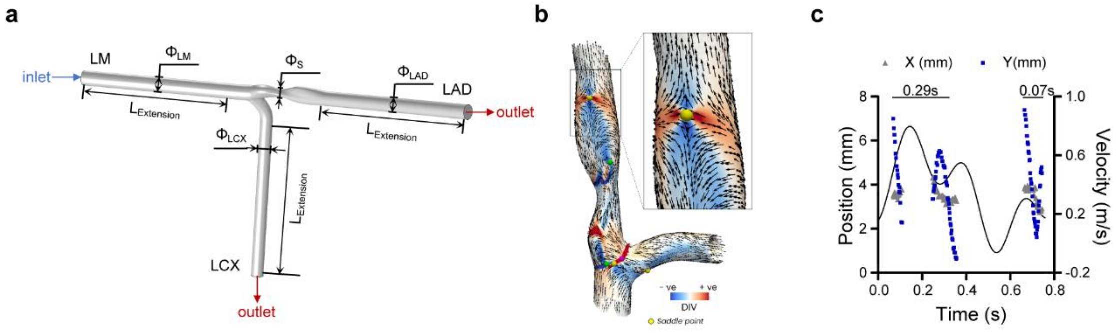


**Fig. S1 | Topological analysis of OTSS in a stenotic coronary artery. a**, LM–LAD–LCX bifurcation geometry reconstructed from patient DSA. The proximal LM is prescribed as a time-varying velocity inlet, and the distal LAD and LCX are set as pressure outlets. **b**, Representative WSS topology on the luminal surface. Black arrows indicate local WSS vectors, and the colormap shows the divergence of the WSS field. The inset highlights a saddle-type fixed point identified downstream of the stenosis, corresponding to a −1 WSS topology core. **c**, Time-resolved trajectories of the core coordinates along the axial (x, gray) and radial (y, blue) directions, plotted together with the inlet velocity waveform on the right axis. Two persistence intervals of the core are detected over one cardiac cycle.

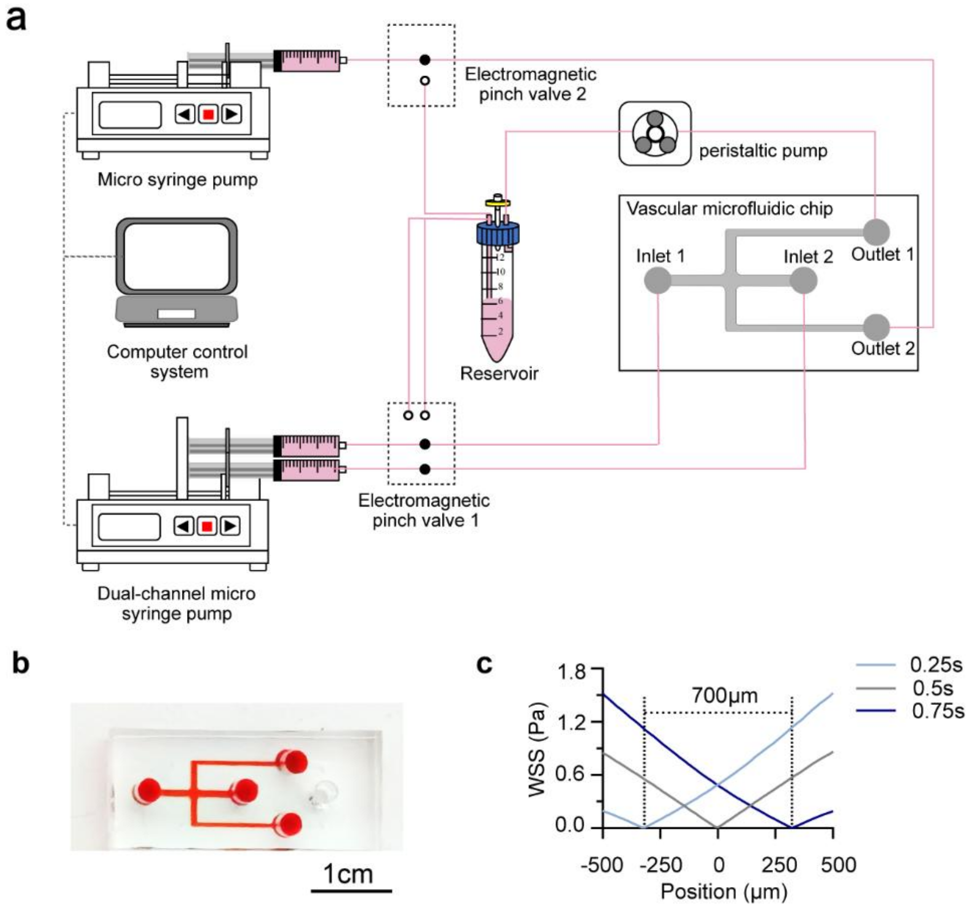


**Fig. S2 | Microfluidic implementation and validation of oscillatory topological shear stress. a**, Schematic of the computer-controlled perfusion system. Two syringe pumps are regulated by a controller to generate the prescribed flow-rate waveforms. **b**, Photograph of the microfluidic chip. **c**, Numerical simulations of WSS profiles along the central axis of the outlet channel at different time points within one oscillation cycle. The oscillation frequency is 1 Hz, and the channel center is defined as zero position. Profile minima indicate instantaneous −1 TSS core positions, with a peak-to-peak displacement of approximately 700 μm.

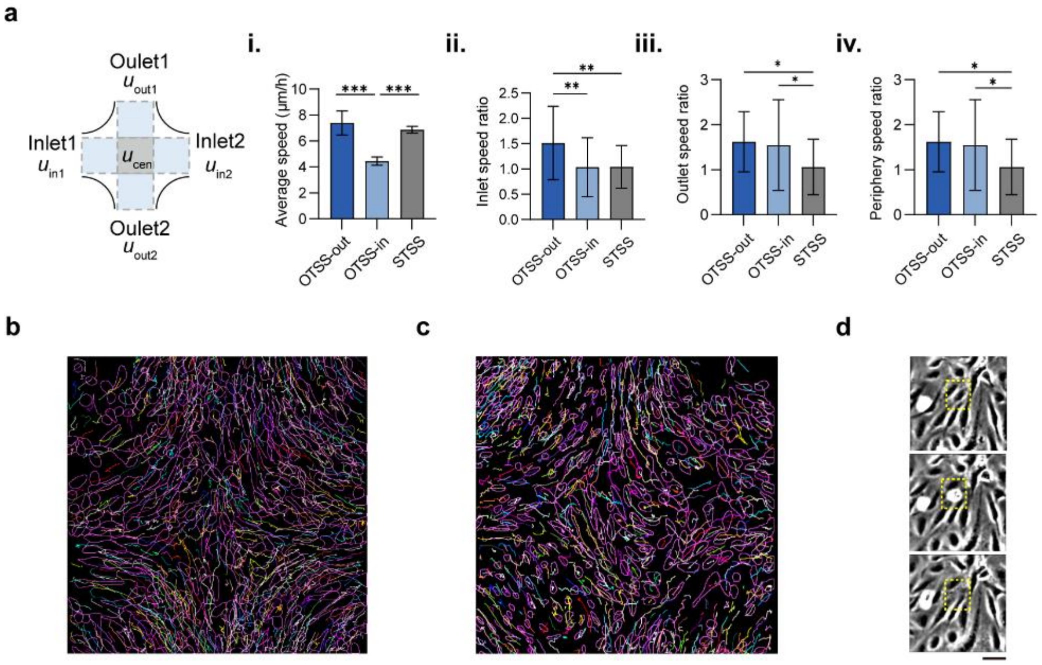


**Fig. S3 | Collective migration, density heterogeneity, and extrusion dynamics of endothelial crowds under different shear conditions.** **a**, Quantification of migration speed under different flow conditions. Schematics delineate channel boundaries: blue regions indicate the two inlet and two outlet zones, and the gray region marks the central −1 TSS region. **i**, Mean velocity in peripheral regions, calculated as the average of area-averaged migration speeds measured in the four straight channel segments (two inlets and two outlets). **ii, iii**, Area-averaged velocities in the paired inlet ($u_{in1,2}$) and outlet ($u_{out1,2}$) regions, respectively. **iv**, Ratio of mean peripheral velocity (average of four channel segments) to area-averaged velocity in the central region. **b**, **c**, Representative single-cell trajectories under OTSS-out (**b**) and STSS (**c**) within a 1000 × 1000 μm field of view. **d**, Time-lapse sequence showing local antiparallel rearrangement and cell extrusion under OTSS. The yellow box marks a representative cell that spreads, bulges apically, and is extruded as neighbors reorganize, illustrating a possible route of density loss in nematic regions. Scale bar, 50 μm.

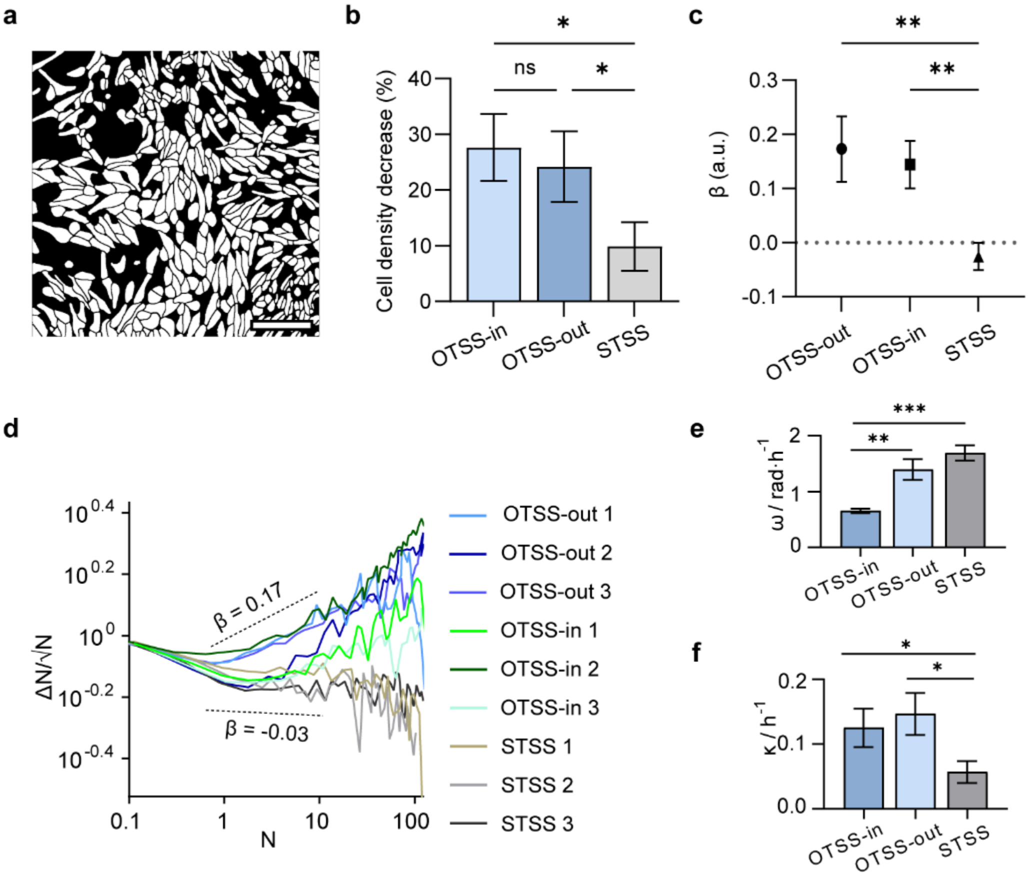


**Fig. S4 | Characterization of endothelial collective dynamics under inlet-axis oscillatory topological shear stress. a**, Representative segmentation mask of an endothelial monolayer exposed to OTSS-in, illustrating pronounced spatial heterogeneity in cell distribution. **b**, Quantification of cell detachment under OTSS-in, OTSS-out and STSS. **c**, Statistical comparison of the fitted scaling exponent β from giant number fluctuation (GNF) analysis across the three flow conditions. **d**, Log–log plots of cell number fluctuations at the end of flow stimulation for three independent experiments under each condition, with dashed reference trend lines corresponding to $\beta = 0.17$ and $\beta = -0.03$. **e**, Fitted oscillation frequency ω extracted from the temporal evolution of normalized defect density under STSS, OTSS-in, and OTSS-out. **f**, Fitted decay coefficient κ obtained from the same analysis.

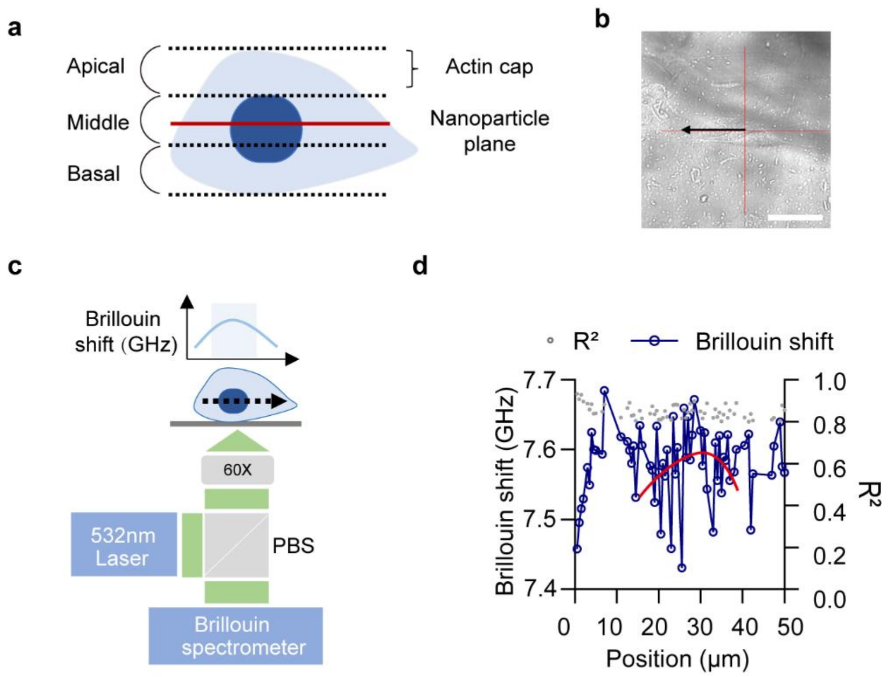


**Fig. S5 | Z-axis-resolved image segmentation and Brillouin microscopy workflows. a,** Schematic of the z-axis partitioning and region definition. Cells are divided into three equal-height z-tiers (apical, middle, and basal). The apical tier is projected and averaged to represent the actin cap. The nuclear mask is eroded to define a perinuclear region used to sample actin cap and NP fluorescence intensity. **b,** Bright-field image showing the scanning pattern across a single cell, with the red cross indicating the initial laser spot position and the black arrow denoting the scanning direction; scale bar, 50 μm. **c,** Schematic of the optical setup for Brillouin microscopy using a 532 nm laser, illustrating point-scanning over the nuclear region, together with representative Brillouin frequency shift spectra. **d,** Representative Brillouin frequency shift spectra fitted with a Lorentzian function ($R^2 > 0.8$), exhibiting a characteristic peak profile within the nuclear region. The mean Brillouin shift for each nucleus is calculated from at least 15 measurement points acquired at 0.5 μm spacing.

**Supplementary Table**

| Symbol | Numerical Value | Unit | Physical Meaning |
|---|---|---|---|
| $\Phi_{LM}$ | 3.6 | mm | The diameter of LM. |
| $\Phi_{LAD}$ | 3.5 | mm | The diameter of LAD. |
| $\Phi_{LCX}$ | 3.0 | mm | The diameter of LCX. |
| $\Phi_{S}$ | 2.5 | mm | The diameter of Stenosis. |
| $L_{Extension}$ | 40 | mm | The length of vascular extension segment. |

**Supplementary Table S1 | Geometric parameters of the stenotic coronary artery model used in numerical simulations.** Characteristic diameters and extension length define the LM–LAD–LCX bifurcation geometry reconstructed from patient DSA.

| Condition | Channel | Flow rate (μL/min) | Notes |
|---|---|---|---|
| STSS | Inlet 1 | $Q_{\mathrm{in},1}(t) = 145$ | Constant flow |
| | Inlet 2 | $Q_{\mathrm{in},2}(t) = 145$ | Constant flow |
| | Outlet 1 | $Q_{\mathrm{out},1}(t) = 145$ | Constant flow |
| | Outlet 2 | $Q_{\mathrm{out},2}(t) = 145$ | Constant flow |
| OTSS-in | Inlet 1 | $Q_{\mathrm{in},1}(t) = 145 + 100\sin(2\pi t)$ | Sinusoidal inflow |
| | Inlet 2 | $Q_{\mathrm{in},2}(t) = 145 - 100\sin(2\pi t)$ | Sinusoidal inflow, phase-shifted by $\pi$ |
| | Outlet 1 | $Q_{\mathrm{out},1}(t) = 145$ | Constant outflow |
| | Outlet 2 | $Q_{\mathrm{out},2}(t) = 145$ | Constant outflow |
| OTSS-out | Inlet 1 | $Q_{\mathrm{in},1}(t) = 145$ | Constant inflow |
| | Inlet 2 | $Q_{\mathrm{in},2}(t) = 145$ | Constant inflow |
| | Outlet 1 | $Q_{\mathrm{out},1}(t) = 145 + 100\sin(2\pi t)$ | Sinusoidal outflow |
| | Outlet 2 | $Q_{\mathrm{out},2}(t) = 145 - 100\sin(2\pi t)$ | Sinusoidal outflow, phase-shifted by $\pi$ |

**Supplementary Table S2 | Flow-rate waveforms used to generate STSS and OTSS conditions in the microfluidic chip.** Inlet and outlet flow-rate waveforms applied to produce stationary and oscillatory topological shear stress conditions, including waveform type, phase relationship and oscillation frequency.

| Condition | Cell loss (%) | $\beta$ (a.u.) |
|---|---|---|
| STSS | 9.9 ± 4.4 | -0.03 ± 0.03 |
| OTSS-in | 27.6 ± 6.0 | 0.14 ± 0.04 |
| OTSS-out | 24.2 ± 6.4 | 0.17 ± 0.06 |

**Supplementary Table S3 | Quantification of cell loss and giant number fluctuations.** Quantitative comparison between experiments and simulations for total cell loss and giant number fluctuation scaling exponent $\beta$ under each flow condition. Experimental values are reported as mean ± s.d..

| **Condition** | **Decay Amplitude, *A*** | **Decay coefficient, *κ* (h⁻¹)** | **Angular frequency, *ω* (rad h⁻¹)** | **Oscillation Amplitude, *B*** | **Phase, *ϕ*** | **R²** |
|---|---|---|---|---|---|---|
| | 0.58 | 0.040 | 1.70 | 0.28 | 1.95 | 0.65 |
| STSS | 0.71 | 0.057 | 1.81 | -0.16 | 0.24 | 0.62 |
| | 0.89 | 0.074 | 1.54 | -0.12 | 4.92 | 0.61 |
| | 0.84 | 0.151 | 1.52 | 0.10 | 5.59 | 0.78 |
| OTSS-out | 0.92 | 0.112 | 1.18 | -0.18 | 0.67 | 0.76 |
| | 0.87 | 0.177 | 1.49 | -0.16 | 7.87 | 0.79 |
| | 0.84 | 0.096 | 0.62 | -0.28 | 2.37 | 0.88 |
| OTSS-in | 0.84 | 0.155 | 0.64 | 0.17 | 2.94 | 0.73 |
| | 0.94 | 0.125 | 0.70 | 0.09 | 3.02 | 0.79 |

**Supplementary Table S4 | Fitting parameters of defect-density dynamics.** Best-fit parameters describing the temporal evolution of normalized ±1/2 defect density under different shear conditions. Each row corresponds to an independent experimental replicate.

| **Condition** | **Diffusion coefficient, $D$ (μm² h⁻¹)** | **Persistence time, $T_p$ (h)** | **$R^2$** |
|---|---|---|---|
| OTSS-in | 76.71 ± 1.25 | 1.46 ± 0.08 | 0.99863 |
| OTSS-out | 148.29 ± 2.93 | 1.74 ± 0.10 | 0.99850 |
| STSS | 187.97 ± 1.76 | 1.74 ± 0.05 | 0.99967 |

**Supplementary Table S5 | Persistent random walk parameters of single-cell migration.** Effective diffusion coefficient and persistence time obtained by fitting single-cell mean squared displacement curves to a persistent random walk model under the indicated flow conditions. Values are reported as mean ± s.d.

**Supplementary Video**

**Supplementary Video 1 | Oscillatory −1 wall shear stress topology in a stenotic coronary artery.** Time-resolved numerical simulation of pulsatile blood flow in a patient-specific stenotic coronary artery. The near-wall wall shear stress (WSS) vector field is shown on the luminal surface, revealing saddle-type fixed points corresponding to −1 WSS topology upstream and downstream of the stenosis.

**Supplementary Video 2 | −1 WSS defect dynamics in microfluidic OTSS-in and OTSS-out chips.** Numerical visualization of wall shear stress (WSS) topology in the cross-shaped microfluidic vascular chip. Sinusoidal modulation of inlet or outlet flow rates induces oscillatory displacement of the −1 TSS core along the inlet (OTSS-in) or outlet (OTSS-out) axis while preserving physiological shear levels in the straight channels.

**Supplementary Video 3 | Time-lapse endothelial crowd dynamics under STSS and OTSS.** Time-lapse imaging of endothelial monolayers exposed to stationary topological shear stress (STSS), outlet-axis oscillatory topological shear stress (OTSS-out) and inlet-axis oscillatory topological shear stress (OTSS-in).

**Supplementary Video 4 | Polar and nematic order in endothelial crowds under OTSS.** Representative time-lapse visualization of endothelial collective organization under oscillatory topological shear stress (OTSS). Cell trajectories and orientations are shown for low-density polar domains exhibiting coherent unidirectional migration and high-density nematic domains characterized by antiparallel streams.
